# Cellular and transcriptional diversity over the course of human lactation

**DOI:** 10.1101/2021.11.13.468496

**Authors:** Sarah K. Nyquist, Patricia Gao, Tessa K. J. Haining, Michael R. Retchin, Yarden Golan Maor, Riley S. Drake, Kellie Kolb, Benjamin E. Mead, Nadav Ahituv, Micaela E. Martinez, Bonnie Berger, Alex K. Shalek, Brittany A. Goods

## Abstract

Human breast milk is a dynamic fluid that contains millions of cells, but their identities and phenotypic properties are poorly understood. We used single-cell RNA-seq (scRNA-seq) to characterize the transcriptomes of cells from human breast milk (hBM) across lactational time from 3 to 632 days postpartum in 15 donors. We find that the majority of cells in human breast milk are lactocytes, a specialized epithelial subset, and cell type frequencies shift over the course of lactation yielding greater epithelial diversity at later points. Analysis of lactocytes reveals a continuum of cell states characterized by transcriptional changes in hormone, growth factor, and milk production related pathways. Generalized additive models suggest that one sub-cluster, LALBA^low^ epithelial cells, increase as a function of time postpartum, daycare attendance, and the use of hormonal birth control. We identify several sub-clusters of macrophages in hBM that are enriched for tolerogenic functions, possibly playing a role in protecting the mammary gland during lactation. Our description of the cellular components of breast milk, their association with maternal-infant dyad metadata and quantification of alterations at the gene and pathways levels provides the first detailed longitudinal picture of human breast milk cells across lactational time. This work paves the way for future investigations of how a potential division of cellular labor and differential hormone regulation might be leveraged therapeutically to support healthy lactation and potentially aid in milk production.

## INTRODUCTION

Human breast milk (hBM) is the nutritional food source evolved specifically to meet the needs of infants.^1^ Feeding exclusively with hBM is currently recommended for the first six months of life, and this is one of the strongest preventative measures against mortality in children under 5 years old.^2^ In addition, breastfeeding has been linked to long-term health benefits for both infants and nursing mothers.^1,3,4^ Breastfed infants have decreased infections^5^, improved gut and intestinal development^6^, and improved regulation of weight long after termination of breastfeeding.^7^ Additionally, nursing mothers have a decreased risk of ovarian and breast cancers.^8–10^ Given that lactation and nursing provide unprecedented health benefits to mothers and infants, there is a need to better understand the molecular and cellular features of hBM, and broadly, how these may correlate with maternal and infant lifestyles and health.

The stages of lactation are canonically described as colostrum (0-5 days postpartum), transitional (6-14 days postpartum), and mature (>15 days postpartum) followed by involution, which begins within hours of the cessation of lactation.^11,12^ During pregnancy, lactation and involution, the human mammary gland undergoes drastic remodeling that requires coordinated shifts in tissue architecture and cellular composition guided by hormonal cues.^13,14^ During lactation, the cells of the mammary gland are responsible for synthesizing and transporting the diverse components of hBM as well as responding to tightly regulated and highly responsive signals maintaining lactational viability. A mechanistic understanding of the cellular composition, activities, and regulation of the human mammary gland in the period between the establishment of lactation and involution is essential for understanding environmental factors that impact milk production, the responsiveness of the breast to the changing nutritional needs of the infant, and the mechanisms of long-term lactation. However, given the unique nature of this tissue niche, it is challenging to study lactating tissue directly in humans.

hBM contains live cells which are thought to enter the breast through exfoliation during the process of breastfeeding, thereby providing an opportunity to study lactational cells.^11,15^ Cells from hBM are viable, can be cultured, and immune cells were shown to transfer to offspring’s bloodstream and tissues in animal models.^12,16–18^ The investigation of these live cells provides both non-invasive surveillance of the cells in the mammary mucosa and allows for a more detailed understanding of their roles in infant development.^12,18–21^ The cellular fraction of hBM contains both somatic and immune cells.^11^ Immune cell populations, such as macrophages^19,22^, may be involved in the protection of the breast itself from infection during lactation^11^. They may also produce important bioactive components, such as antibodies and cytokines, which play a role in the establishment of the infant immune system.^23^ Somatic cells identified in breast milk include epithelial cells and a small fraction of stem cells.^11^ Studies have identified both ductal myoepithelial cells and secretory epithelial cells (lactocytes) in breast milk, where the latter predominates.^11^ Lactocytes are involved in the synthesis and transport of an array of factors, such as human milk oligosacharides (HMOs), lactose, micronutrients, fat, hormones, cytokines, into the lumen of the lactating breast. Much remains to be learned about the mechanisms by which these essential components are created and transported into breast milk and how the behaviors of these cells are regulated.^11,13,24^ Despite their dual role in producing dynamic nutrition for infants and conferring immunological protection, it is still unclear how they may change over the course of lactation.^3,4^

To date, several studies have used either bulk^12,25–28^ or single-cell RNA-sequencing (scRNA-seq)^16,29^ to study the transcriptome of hBM in small cohorts. These studies have revealed subsets of epithelial cells in hBM, as well as progenitor luminal cells, and genes that change in bulk over the course of lactation. Bulk analysis, however, limits our ability to delineate key cell states and uncover specialized cell phenotypes.^30,31^ scRNA-seq analyses to date have also been limited by low sample numbers and small donor pools, thereby decreasing the ability to characterize the cross-donor heterogeneity of breast milk longintudinally.^1,28,32^ Longitudinal studies of other factors in milk composition have characterized dynamic shifts in hormone concentrations^33–35^, cytokine content^36–38^, and overall protein content^39^ up to 3 months postpartum suggesting that most components decrease in concentration early in lactation. However, no transcriptomic studies to date have captured the full range of lactation across time. How these dynamic milk changes relate to lactocytes and immune cell function, are also not well understood.^11^

In order to better understand cellular dynamics and longitudinal lactational heterogeneity, we sought to characterize the transcriptomics of hBM-derived cells using scRNA-seq on longitudinal samples. hBM was collected longitudinally from 15 human donors across various stages of lactation (Supplemental Table 1, Figure 1A). For each sample, we collected a rich set of information about the mother-infant dyad, including vaccine history, illness, and daycare status. To our knowledge, we have generated the first single-cell analysis of hBM-resident cells over the course of lactation, with a dataset comprised of over 48,478 cells from 50 samples. We identify key cell subsets, including immune cells and epithelial cells at each lactation stage. We further identify several factors that are associated with alterations in cell frequencies over lactation, including the use of hormonal birth control and the start of daycare. We also nominate many pathways and genes that are altered in epithelial subsets over the course of lactation, including those that may be hormonally regulated. Taken together, our data provide the first longitudinal characterization of single cells in breast milk and shed light on the gene programs that may drive crucial human lactocyte functions over the course of lactation.

**Figure 1:**
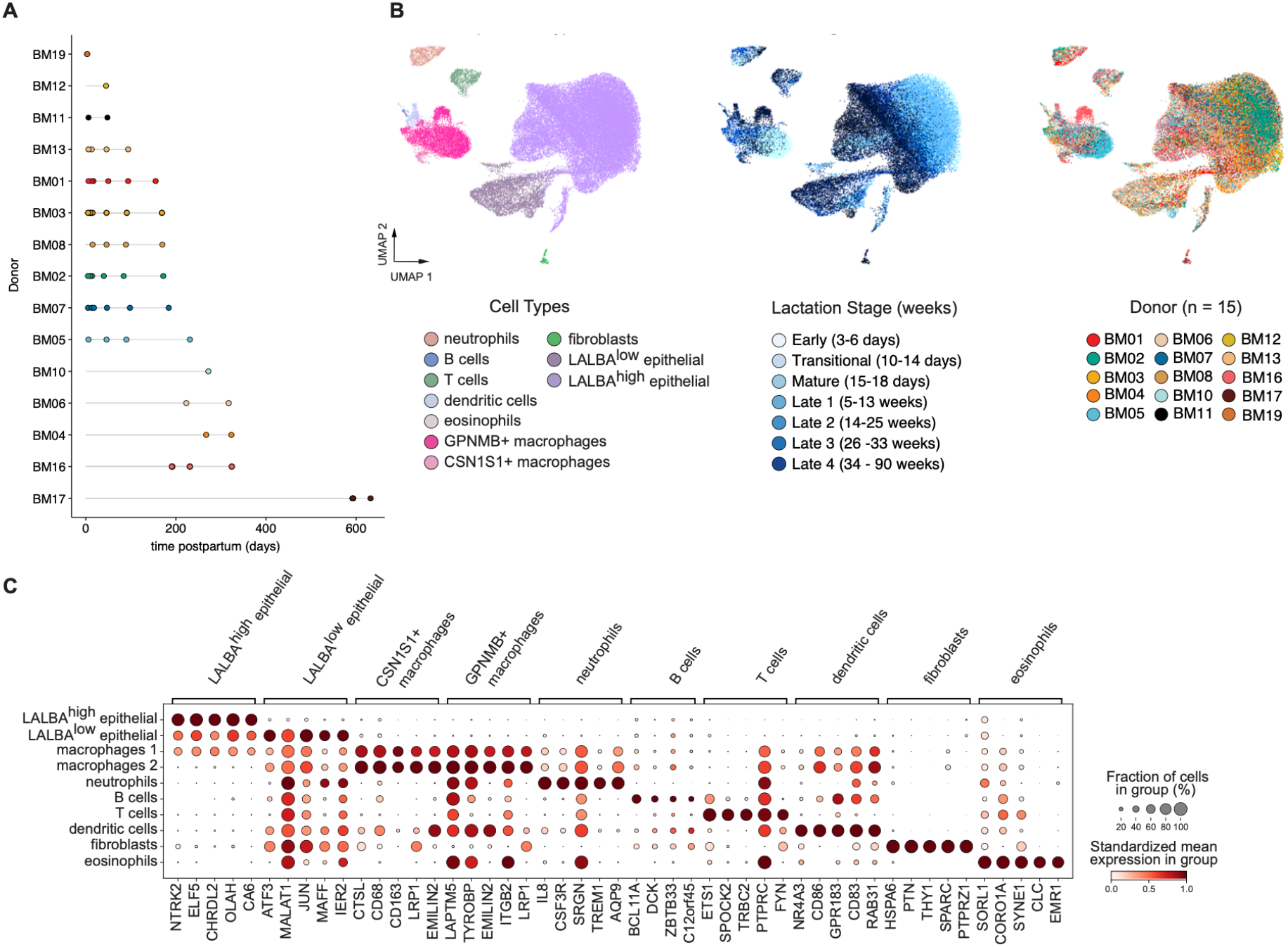
Atlas of cell types present in human breast milk across lactation. **A**. Sampling timeline showing collection of samples for each donor as a function of time postpartum (days). **B**. Projection of dimensionality reduced (Uniform Manifold Approximation and Projection (UMAP)) scRNA-seq data (n = 48,478 cells across 15 donors) colored by cell type, lactation stage (early, transitional, mature, and several late stages), and donor. **C**. Marker genes (x axis) for each major cell type cluster (y axis). Circle size describes percent of cells in cluster expressing the gene. Color represents the mean log-normalized gene expression in that cluster standardized across clusters within each gene.

## METHODS

### Donor enrollment and breast milk collection

Donors were enrolled in the MIT Milk Study under an approved protocol (Protocol # 1811606982). Donors were recruited at hospitals, research institutes, and clinics around the Boston, MA, USA area primarily on the MIT campus. Donors expressed milk using their method of choice and, where possible, provided that information in questionnaires for each sample. To minimize diurnal variations in cell composition, donors provided milk in the mornings between 6:00AM-9:00AM.^40,41^ We also collected extensive donor-supplied metadata for each sample (Supplemental Table 8), including information about maternal and infant health. Donors collected a minimum of 0.5mL of milk, placed in study-provided sample collection bags, and kept on ice until the sample was collected. Samples were processed as close to expression as possible (up to 6 hours) and kept on ice until cells were isolated. Donors also provided answers to the study questionnaire with each sample. Donors provided milk at various time points, covering the following milk stages: early 3-5 days postpartum (colostrum/early), transitional (10-14 days), mature (15-18 days), and several later stages (late 1: 5-13 weeks, late 2: 14-25 weeks, late 3: 26-33, and Late 4: 34-90 weeks). Breast milk was sampled from 15 mothers between the ages of 25-34 (median age 31). All pregnancies were full term with seven donors reporting induced labor, four reporting C-sections, and all but two donors reporting no prior pregnancies. Four donors began hormonal birth control during the sampling period. Eight total samples from six donors were collected after starting day care.

### Cell isolation

To isolate cells directly from whole milk, samples were processed as previously described.^42^ Briefly, milk was diluted 1:1 with cold PBS and cells were pelleted by centrifugation for 10 minutes at 350g. After removal of skim milk and the fat layer, cells were transferred to a clean tube in 1mL of cold PBS and washed three times in 10 mL of cold PBS. The final cell pellet was resuspended in 1mL of cold complete RPMI media (ThermoFisher) containing 10% FBS and 5% pen/strep (ThermoFisher). Cells were counted with a hemocytometer and Seq-Well S^3^ was performed as described below^43^. For experiments comparing milk handling and cell isolation methods, cells were isolated as described above from milk that had been sorted at 4°C or at -20°C overnight. Frozen milk was thawed in a 37°C water bath prior to cell isolation. For sorting of live cells, milk cells were isolated directly from milk and stained according to the manufacturers protocol for calcein violet (ThermoFisher) and sytox green (Invitrogen) prior to sorting for calcein violet positive and sytox green negative cells on a Sony Sorter (SH800S). For enrichment of live cells, directly isolated milk cells were processed according to the manufacturer’s instructions (EasySep Dead Cell Removal (Annexin V) Kit).

### Generation of single-cell RNA-sequencing (scRNA-seq) data with Seq-Well S^3^

Seq-Well S^3^ was performed as described previously.^43,44^ For each milk sample, about 15,000 cells were loaded onto each array preloaded with uniquely-barcoded mRNA capture beads (ChemGenes). Arrays were washed with protein-free RPMI media, then sealed with polycarbonate membranes. Arrays were incubated at 37°C for 30 minutes to allow membranes to seal, then transferred through a series of buffer exchanges to allow for cell lysis, transcript hybridization, bead washing, and bead recovery from arrays post membrane removal. Reverse transcription was performed with Maxima H Minus Reverse Transcriptase (ThermoFisher), excess primers were removed using an Exonuclease I digestion (New England Biolabs), second strand synthesis was performed, and whole transcriptome amplification (WTA) by PCR was performed using KAPA Hifi PCR Mastermix (Kapa Biosystems). WTA product was purified using Agencourt Ampure beads (Beckman Coulter) and dual-indexed 3’ digital gene expression (DGE) sequencing libraries were prepared using Nextera XT (Illumina). Libraries were sequenced on a NovaseqS4 or NovaseqS2 with a paired end read structure (R1: 20 bases; I1: 8 bases; I2: 8 bases; R2: 50 bases) and custom sequencing primers.

### Analysis of scRNA-seq data

#### Alignment and quality control

Data was aligned using the Dropseq-tools pipeline on Terra (app.terra.bio) to human reference genome hg19. Sequencing saturation curves were generated using custom scripts to ensure adequate sequencing depth (data not shown).

#### Clustering and cell identification

Samples were split into milk stage groups for initial clustering and doublet identification. For each sample, scrublet was run with default parameters and cells identified as doublets were removed from downstream analysis.^45^ For each milk stage, all samples were combined into a single scanpy object, cells were filtered with parameters: >400 genes, >750 UMI, <750 counts, <20% UMIs from mitochondrial genes. UMI counts were log-normalized and the top 2000 variable genes were identified with the batch_key parameter set to “sample”. PCA was run on scaled data, and a nearest neighbors map was calculated with 15 neighbors and 25 principal components prior to running UMAP for visualization. Resulting clusters were robust to multiple choices of clustering parameters. Clustering of resulting DGEs was performed using Leiden clustering in the Scanpy (scanpy.readthedocs.io) package independently on samples of each milk stage.^46^ Clusters were classified as immune cells or epithelial cells for further sub-clustering based on expression of *PTRPC* (immune cells) and *LALBA* (epithelial cells). Upon sub-clustering on each of these subsets, doublets were identified as clusters co-expressing multiple lineage markers and were removed. Sub-clustering was performed on the applicable clusters from all time points combined.

#### Pseudobulk marker gene identification

To identify marker genes for celltype clusters whose specificity to Leiden clusters or cell subgroups was consistent across donors and samples, we utilized pseudobulk marker gene identification^47–49^. Raw gene expression counts were pooled by sample and cluster such that one pseudobulk population was created for each cluster found in each sample. Psuedobulk groups were filtered to include only sample-subcluster pairs containing at least 10 cells. Differential expression between clusters of one celltype and all other clusters was executed using a Wald test in DESeq2^50^ with the design formula “∼donor + is.thiscelltype” where the factor ‘is.thiscelltype’ is set to TRUE for pseudobulk populations from the cluster of interest and FALSE for other clusters. These pseudobulk marker genes were filtered for adjusted p value < 0.05, percent expression of single cells in the cluster > 30%, and DESeq2-calculated log2 fold change > 0.4. Pseudobulk marker genes of all cell types (Supplemental Table 2) and epithelial cell groups (Supplemental Table 3) and top marker genes sorted by difference in percent of cells expressing in-cluster compared to out-of-cluster are visualized in Figure S7E and Figure 3D, respectively.

#### Epithelial cell sub-clustering

Epithelial sub-clustering was performed on combined cells from all samples to identify major cell states within the data and characterize their changes in gene expression over the course of lactation. To enable these analyses, we identified cell groups which were either distinct enough to be robust to clustering parameter selection, or, for groups of cells whose core identifying gene expression profiles could not be defined with respect to other clusters, similar clusters were merged and further analysis identified genes changing over time. Sub-clustering proceeded by re-discovering the top 3,000 variable genes on the epithelial subset, re-running PCA on these genes, and clustering with Leiden clustering with resolution 0.5 and 10 neighbors on 22 principal components (Figure S4A). Clusters 0, 1, 2, and 3 were merged into the secretory lactocyte cluster due to shared expression of various canonical lactation-related genes (Figure S7F). Despite many shared functions with clusters 0, 1, 2, and 3, cluster 5 was left as its own cluster due to high mitochondrial gene percentage (Figure S7G). Clusters 9, 6 and 8 shared a distinct transcriptional signature and were merged into the LALBA^low^ epithelial cluster. Clusters 4 and 11 were merged into a single KRT high 1 cluster due to cluster 11’s specificity to a. single donor, and cluster 7 remained as a single KRT high 2 cluster. Additionally, these clusters were robust to leave one out clustering.

#### Immune cell sub-clustering

Immune cells were sub-clustered separately and re-filtered to remove additional doublets. To accomplish the latter, immune cells were clustered with a known subset of secretory epithelial cells from our epithelial cell data. This allowed us to generate a gene signature derived of PC1-specific genes to define lactocytes or monocytes with high confidence (Supplemental Table 4). We performed module scoring with these in R (v3.6.2) with Seurat (V3), allowing us to stringently filter for immune cells that scored highly for lactocyte gene expression (>2.5 standard deviations above the mean lactocyte module score)^51^. Finally, we identified any additional doublets based on dual expression of key lineage markers as described above. We performed sub-clustering analyses by re-normalizing the data, finding the top 2000 variable genes, re-scaling the data, running PCA, then performing additional UMAP visualization with the first 15 principal components. Supervised marker gene identification was performed across cell types using Seurat’s Wilcoxon rank-sum test. We also performed sub-clustering analyses on the monocytes and macrophages as these were the most abundant immune cell type. These cells were re-normalized, the top 2,000 variable genes were identified, and the data was clustered across several resolutions to identify resolutions that produced non-redundant clusters (resolution = 0.2) as determined by marker gene identification using Seurat’s Wilcoxon rank-sum test.

#### Identification of time-varying genes

Time-associated genes were identified for each cluster using pseudo-bulk analysis. First the raw counts all cells in each sample in each cluster were summed to create sample and cluster specific pseudobulk data. Then DESeq2 was used to identify genes varying over the course of lactation in each sub-cluster using a likelihood ratio test between the design formula “∼ 0 + donor + days_post_partum” over “∼0 + donor”. Samples with a minimum of 10 cells in a cluster were included in the analysis, and samples from more than 400 days postpartum were excluded from time series analyses to avoid the small number of very late samples driving a disproportionate amount of variation due to the large gap in time between samples before 400 days postpartum and after. Genes with in-cluster single cell percent expression > 20% and adjusted p value <0.05 were included in downstream visualization and enrichment analyses. Heatmaps represent row-z-scored, log normalized per-sample expression of genes of interest. Principal component analysis on pseudobulk samples from each epithelial subset was used to identify the primary axis of variation within each subset by identifying the sample metadata and genes correlated with the first principal component. The first principal component of the LALBA^low^ epithelial and secretory lactocyte subsets was highly correlated with time postpartum, so time dependent gene analyses were focused on these subsets (Supplemental Table 5A,B). We classified universal epithelial cell time varying genes as genes associated with time and changing in the same direction in both LALBA^low^ epithelial and secretory epithelial subsets (Supplemental Table 5C,D). Time varying genes in opposite directions in the LALBA^low^ epithelial and secretory epithelial subsets were also identified (Supplemental Table 5E).

#### Identification of metadata associated cellular populations

Associations between collected covariates and cellular population proportions were tested using generalized additive models. For each sample, cell cluster proportions were calculated from the numbers of cells found in each broad celltype by dividing the number of cells in that cluster by the total cells in that sample. Then a generalized additive model was run for each celltype on samples collected earlier than 400 days postpartum using the mgcv R package with model formula ‘celltype_proportion ∼ donor + s(time_post_partum_days, k=7)’.^52^ Additional covariates – including: daycare attendance, infant illness, breast soreness, supplementation with formula, use of hormonal birth control, solid food consumption, and recent vaccinations were tested with model formulas following the pattern ‘celltype_proportion ∼ donor + <covariate> + s(time_post_partum_days, k=7)’. Only samples with complete metadata for a given covariate were included in the corresponding comparison (Supplemental Table 6). In cases where multiple covariates were significantly associated with one celltype proportion, a model including both was run. Specifically, LALBA^low^ epithelial cell proportion was modeled as ‘LALBA^low^ _proportion ∼ donor + daycare + hormonal_birthcontrol + s(time_post_partum_days, k=7)’. Full model results are shown in Supplemental Table 6.

#### Functional enrichment analysis on epithelial cells

Functional enrichment analysis on top marker genes was performed using Enrichr using the gseapy package with the gene set GO_Biological_Processes_2021.^53,54^ Due to the hierarchical structure of the GO database and the overlapping functions of many of the marker genes of the epithelial cell subclusters, representative GO terms were identified through a series of filtering and curating steps. For each subcluster, significantly enriched terms were grouped based on shared marker genes found to be overlapping with the GO term. These grouped terms were further grouped between subclusters based on shared term ID or shared genes. The mean gene set score was calculated for each epithelial cell group and enriched GO term using the scanpy function “score_genes”. For each group of GO terms, the terms with the highest variance of mean gene scores across epithelial subgroup was chosen such that each epithelial subgroup had between 7 and 15 GO terms for which they had the maximum mean gene score. To avoid redundant terms, GO terms were also merged based on high overlap of genes in the full reference GO term gene list. Heatmap visualizations display per-subset mean gene set score for all genes in the GO term z-scored across subsets. Time-dependent enriched GO terms were identified for genes positively and negatively associated with time postpartum separately and for both LALBA^low^ epithelial and secretory lactocyte clusters. These GO terms were similarly curated with an additional filtering step of correlation of the gene set scores over time postpartum in the same direction as the set of differential genes used (e.g. positive correlation for GO terms enriched in the gene list increasing with time). GO terms identified to be changing in the same direction in both the LALBA^low^ epithelial and secretory lactocyte clusters were considered epithelial cell-wide time-varying processes.

#### Statistics

In order to determine if cell fraction in hBM correlated with time postpartum in a subset of continuously sampled donors, we performed a Spearman correlation analysis in R (v4.0.4) using the ggpubr package (v0.4.0). Spearman rank coefficients and associated p values were calculated and displayed, along with confidence intervals, for each cell type over time.

#### Data and code availability

Notebooks to reproduce all analyses performed in R and Python are for download (https://github.com/ShalekLab). Sequencing data are available for download as part of The Alexandria Project (https://singlecell.broadinstitute.org/single_cell?scpbr=the-alexandria-project) and on the Gene Expression Omnibus (GEO ######).

## RESULTS

### We identify major cell types of the breast epithelium and immune cells in human breast milk over the course of lactation

We first optimized a process for generating scRNA-seq data from cells in hBM. Previous studies characterize how sample handling, as well as methods used for cell isolation, can significantly impact the transcriptomes of isolated cells.^55,56^ We compared several workflows for upstream handing of collected hBM – including: fresh isolation of cells, holding at 4°C overnight until cell isolation, and a single freeze thaw of whole milk before isolating cells, as well as several methods for isolating cells, including: sorting live cells, live cell enrichment with a bead-based kit, or centrifugal isolation of fresh cells as previously described (Supplemental Figure 1). We found that for each method, except for freezing, quality control metrics were comparable and we identified expected cell types in milk, including epithelial and immune cell subsets (Supplemental Figure 1B and C). Fresh processing, sorted cells, or live-enriched cells clustered together in PC space, suggesting little gain by additional processing prior to performing scRNA-seq. Additionally, we found that in one donor, fresh but not frozen processing allowed us to retain macrophages (Supplemental Figure 1D). In agreement with previous studies, we found that isolation of cells from fresh milk resulted in the highest quality data and we therefor used this method for our samples analysis.

To better understand the transcriptomes of single cells in hBM over the course of lactation, we recruited donors to provide milk samples at several time points postpartum, including colostrum/early (3-6 days), transitional (10-14 days), mature (15-18 days), and several late points postpartum (5-90 weeks) (Figure 1A). We performed Seq-Well S^3^ with freshly isolated cells from whole milk to generate high quality single cell transcriptomic data across all lactation stages (Supplemental Figure 2).

We performed unsupervised clustering across all high-quality cells and identified cell types using previously identified marker genes (Supplemental Table 2) in the context of the mammary gland and the immune system^57–59^. Our analyses revealed 10 broad cell types representing both epithelial and immune cell compartments (Figure 1B). We identified seven top-level immune cell clusters, including B cells (*TCF4, SEL1L3, CCDC50*), dendritic cells (*NR4A3, REL*), T cells (*ETS1*), two macrophage clusters (GPNMB+ macrophages (*CD68, GPNMB, CTSL*) and CSN1S1+ macrophages (*CD68, CSN1S1, XDH*)), neutrophils (*IL8, CSF3R*), and eosinophils (*SORL1, CORO1A*). We also identified three non-immune top-level clusters, including LALBA^high^ epithelial cells (*XDH, CSN1S1, CSN3*), LALBA^low^ epithelial cells (*CLDN4, JUN, KLF6*), and fibroblasts (*SERPINH1, PTN*). These subsets agree with other datasets describing scRNA-seq on hBM in smaller cohorts.^16,29,32^ We did not identify any basal epithelial cells (Supplemental Figure 2), consistent with previous reports^16,29^. Interestingly, we found that our data clustered predominantly by cell type, rather than donor, suggesting that donor-to-donor differences were not the primary axis of variation. Overall, lactocyte epithelial cells (LALBA^low^ and LALBA^high^) were the most abundant cell type across both donor and lactation stage (mean 81.7% of all cells per sample, standard deviation 24%), with macrophages comprising the most abundant immune cell type (50.5% of immune cells per sample, standard deviation 34%) (Figure 2A).

### Cell frequencies are dynamic over the coure of lactation and associate with maternal-infant metadata

In order to better understand the longitudinal variation in hBM-derived cells, and the overall composition of our cohort metadata, we plotted total cell counts and cell type frequencies over time for each sample in our cohort (Figure 2A). We found that the total cell counts per milliliter of milk decreased over the course of lactation, agreeing with previous literature showing a decrease in total cell counts in mature milk (Supplemental Figure 3).^25^ We also found that the majority of our cohort were directly breastfeeding, with 5 donors (9 samples) additionally supplementing with formula and six donors (9 samples) reporting supplementation with solid foods. Several donors reported breast soreness periodically over the course of the study, with only one donor reporting mastitis at sample collection (Supplemental Table 7). Additionally, none of our donors reported menstruating at the time of sample donation and four were on hormonal birth controls or other reported medications. Finally, we had three donors that had begun weaning and six whose children had started daycare during our study. Globally, the variability in reported metadata allowed us to determine how cellular composition may be impacted by shifts in time, lifestyle and maternal and/or infant health status.

**Figure 1:**
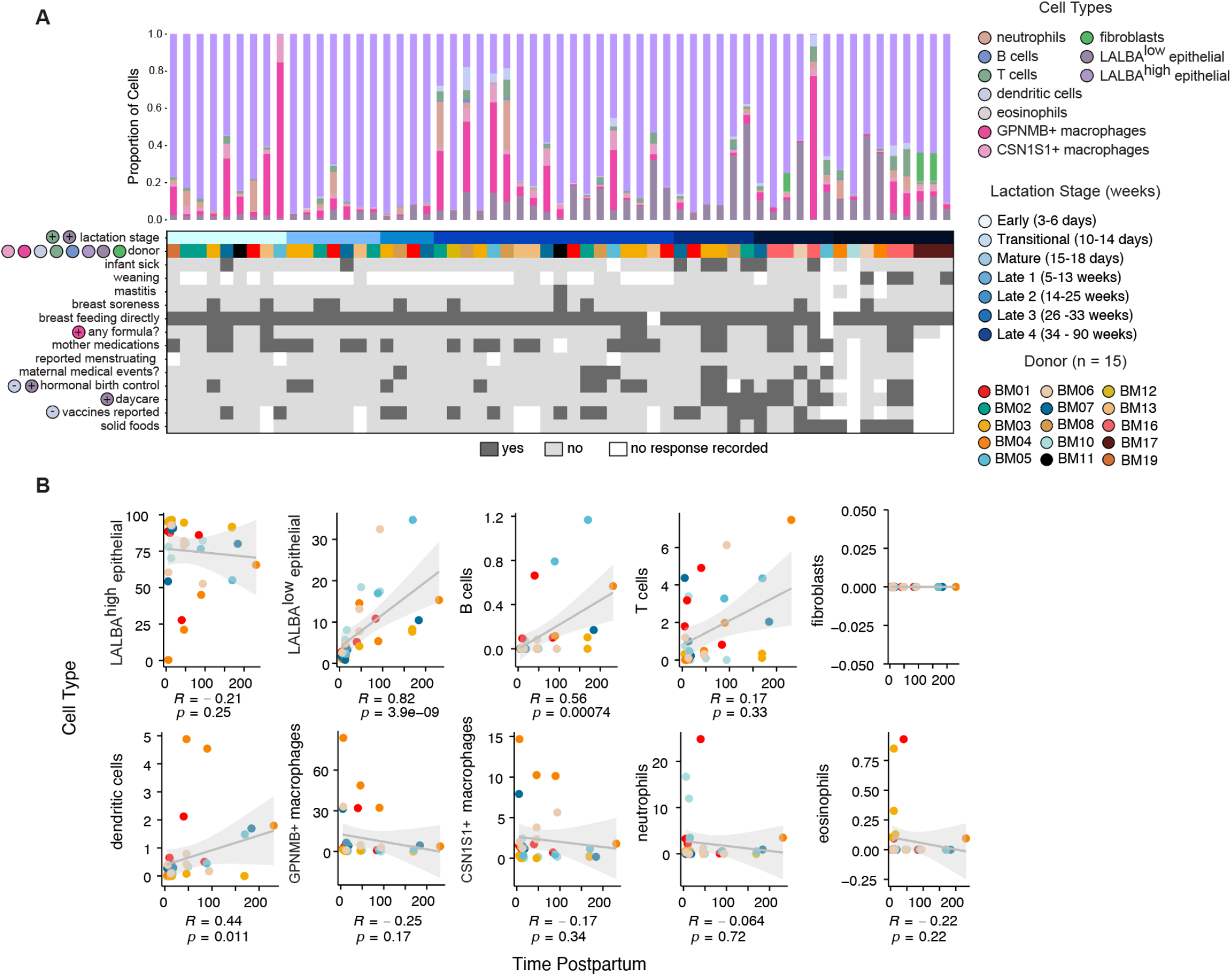
Frequency of cell types over the course of lactation. **A**. Frequency of cell types identified for each sample (top) and associated maternal and infant health information metadata (bottom) collected in user-reported questionnaires. Colored circles to the left of metadata names indicate associations of metadata with cell type (specified by color) abundances and the direction of the association via + or -. Different donors show associations in different directions with celltype’s proportions (see Supplemental Table 6B) **B**. Normalized cell frequencies as function of time postpartum for donors that provided at least three samples are shown for all identified cell types. Spearman correlation coefficients (R) and p values are shown below each plot, and confidence intervals are displayed in grey.

We tested the association between the abundance of identified cell types with any reported metadata using generalized additive models (Supplemental Table 6). While we found that nothing was significantly associated following correction for multiple hypotheses, we did find some associations indicating potential heterogeneity. We found that macrophage 1 proportion associated with formula supplementation, LALBA^low^ epithelial cell proportion positively associated with daycare attendance and with use of hormonal birth control, and dendritic cell proportion negatively associated with use of hormonal birth control and with infant vaccinations (Figure 2A). We noted that a substantial amount of variability in these cell compositions can be attributed to individual donors with a single donor consistently showing substantially larger macrophage proportions (BM05) and all of the fibroblast cells coming from two donors (BM16, BM17). We acknowledge that given our study design, often donor is conflated with certain metadata features.

We next sought to refine our understanding of which cell types were correlated with time postpartum by looking at a subset of donors with at least three samples collected over the course of the study. We found that several cell types remained relatively consistent over the sampled course of lactation, including LALBA^high^ epithelial cells and macrophages (Figure 2B). We also found several cell types that were significantly positively correlated with time postpartum, including LALBA^low^ epithelial cells (p = 2.9e-9) and B cells (p=0.00074) (Figure 2B). Generalized additive models including all samples from fewer than 400 days postpartum also identified LALBA^low^ epithelial cell proportion as positively associated with time postpartum (Supplemental Table 6). Alterations in the composition of the epithelial compartment suggest some emergent cellular functions that support later lactation, and the presence of more B cells or T cells, while still very low fractions of total immune cells, could reflect increasing infant or maternal illnesses reported at later time points in our cohort.

### Macrophages in human breast milk have unique transcriptional and functional programs

We found that the majority of immune cells in hBM over the course of lactation were macrophages, agreeing with previous literature.^60^ We next wanted to better understand the potential functions and phenotypes of macrophages in hBM given that their percentages were altered in response to formula supplementation. We performed sub-clustering analyses and functional enrichment of marker genes that were identified for each sub-cluster (Figure 3A and 3B, Supplemental Table 8). We found five sub-clusters of macrophages that span lactation stage, where macrophage sub-cluster 0 is predominantly from early milk stages, and macrophage sub-cluster 3 is predominantly from later stages and donor BM16. Macrophage sub-clusters were defined by distinct gene signatures and pathway enrichment results (Figure 3B and C). Macrophage sub-clusters 0, 1, and 4 were defined by pathways related to interactions with T cells, neutrophils, and immune tolerance, including IL-10 and PD-1 related pathways. These enrichments were driven by unique sets of genes present in each sub-cluster (Supplemental Table 11). Interestingly, macrophage sub-cluster 0 was defined by several marker genes characteristic of lipid-associated macrophages (*LIPA, TREM2*) and those involved in iron regulation (*FTL*).^61^ Macrophage sub-cluster 3 was enriched for several translation-related pathways, and defined by lipid-related genes like *SCD* and *LTA4H*, and stress-response genes like *NUPR1*. We caution that this sub-cluster was predominantly comprised of one donor, BM16, and thus may reflect specific variations in myeloid cell state related to that particular donor and time point during lactation. Finally, macrophage sub-cluster 2, which was comprised almost entirely of milk macrophages, was defined by structural pathways, transport, and keratinization. This may suggest that these macrophages are important for structural maintenance or have altered their transcriptional state in response to their local tissue milieu, possibly via phagocytosis.^62^ Future work should explore these mechanisms since hBM components have been shown to promote tolerogenic phenotypes in myeloid cells.^63,64^

**Figure 3:**
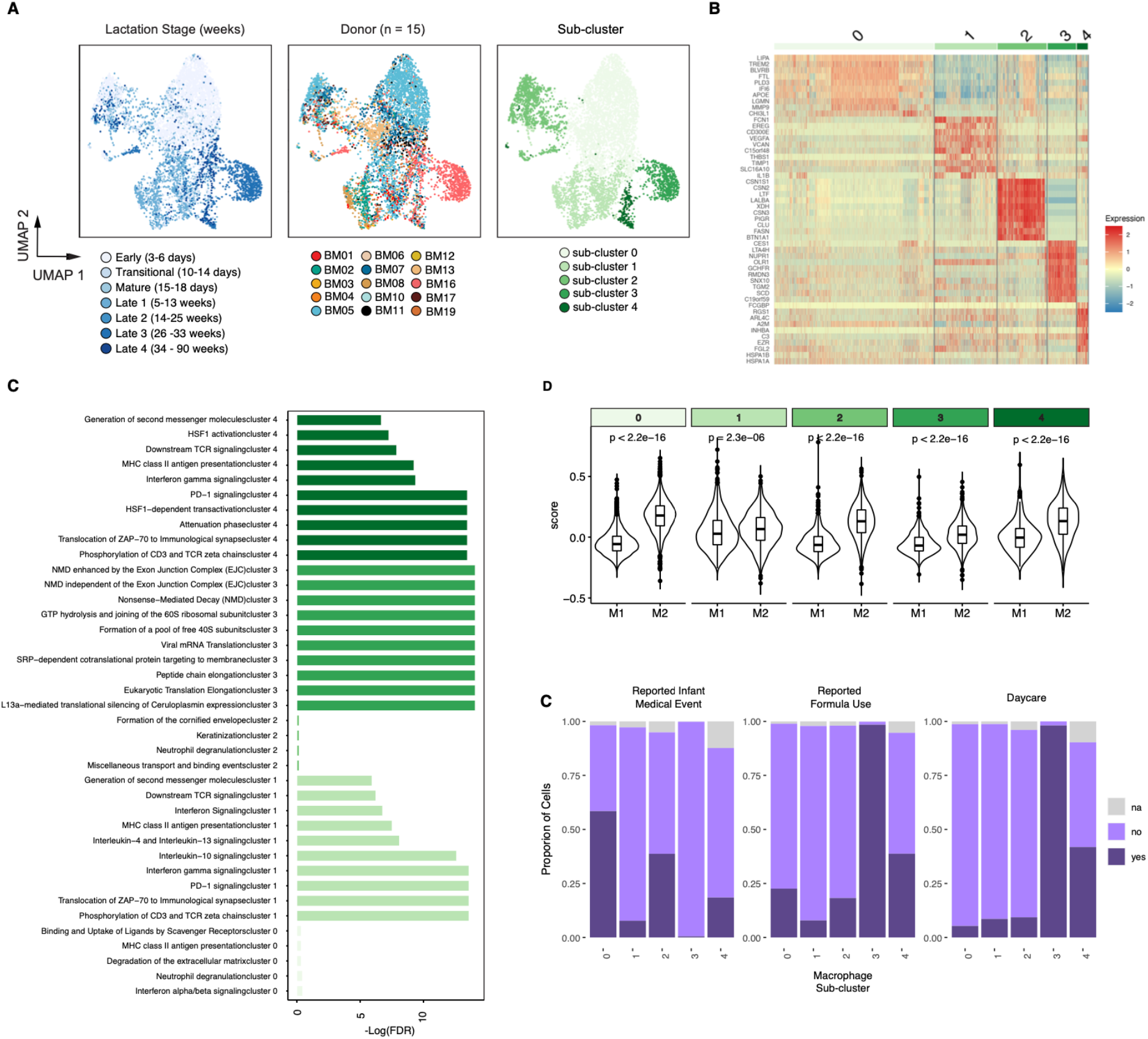
Macrophage sub-clusters across lactation stage. **A**. UMAP projection of hBM-macrophages, colored by lactation stage (left), donor (middle), and macrophage sub-cluster (right). **B**. Heatmap of top marker genes for each identified macrophage sub-cluster. **C**. Reactome enrichment results for each sub-cluster. Full results are shown in Supplemental Table 9. **D**. Module scoring results for M1 or M2 gene sets for each sub-cluster. **E**. Composition of each sub-cluster as a function of infant medical events, weaning status, and daycare.

In order to determine if macrophages in each cluster were more inflammatory (M1) or anti-inflammatory (M2) in nature, we scored these clusters for M1 or M2 gene signatures.^61,65^ While it is widely recognized that macrophages adopt a diverse array of phenotypes in the context of tissues, conventional M1 or M2 status is a useful indicator and comparison point to existing literature in the context of the lactating mammary gland.^19,58^ To accomplish this, we generated module scores for M1 or M2 gene sets within each macrophage sub-cluster. Overall, each sub-cluster, except for sub-cluster 1, scored higher for M2-gene sets, suggesting the majority of macrophages in hBM are M2-like (Figure 3D). Combined with our enrichment results, and previous literature reports in the context of the mammary gland, this suggests that macrophages in hBM predominantly serve immunosuppressive and tissue maintenance functions^19,66^.

Finally, we determined if three meta-data variables of interest, including infant medical events, weaning status, and daycare status, had any compositional variation across sub-clusters (Figure 3E). We found that sub-cluster 0 had the highest proportion of reported medical events, which includes both vaccines and illness. Second, we found that weaning-derived macrophages were predominantly found in sub-cluster 4 (Figure 3E). Future work should address the functional changes in macrophages in hBM post-weaning, since it is known that macrophages shift their transcriptional and functional phenotypes dramatically in response to alterations in the mammary gland.^19^

### Epithelial cell sub-clusters in hBM are enriched for distinct functions and diversify over the course of lactation

In order to better understand the full heterogeneity of epithelial cells in hBM over the course of lactation, we performed sub-clustering analysis on the epithelial cells (see Methods). We identified six sub-clusters of epithelial cells (Figure 4A, Supplemental Figure 4). We found that all epithelial sub-clusters expressed genes related to milk synthesis, such as *LALBA, CSN2, XDH*,^12^ and *FASN* as well as canonical luminal cell markers (*EPCAM, KRT18, KRT19)*, suggesting a clear luminal lineage and role in milk production (Figure 4B)^12^. We also found that there was heterogeneous expression of several canonical mature mammary luminal markers (*KRT18, KRT19*)^12^, hormone receptors (*PRLR, INSR*, and *ESR1*), and stem cell markers (*SOX9, ITGA6*) that have previously been studied in the context of hBM-derived cells^42^.

**Figure 4:**
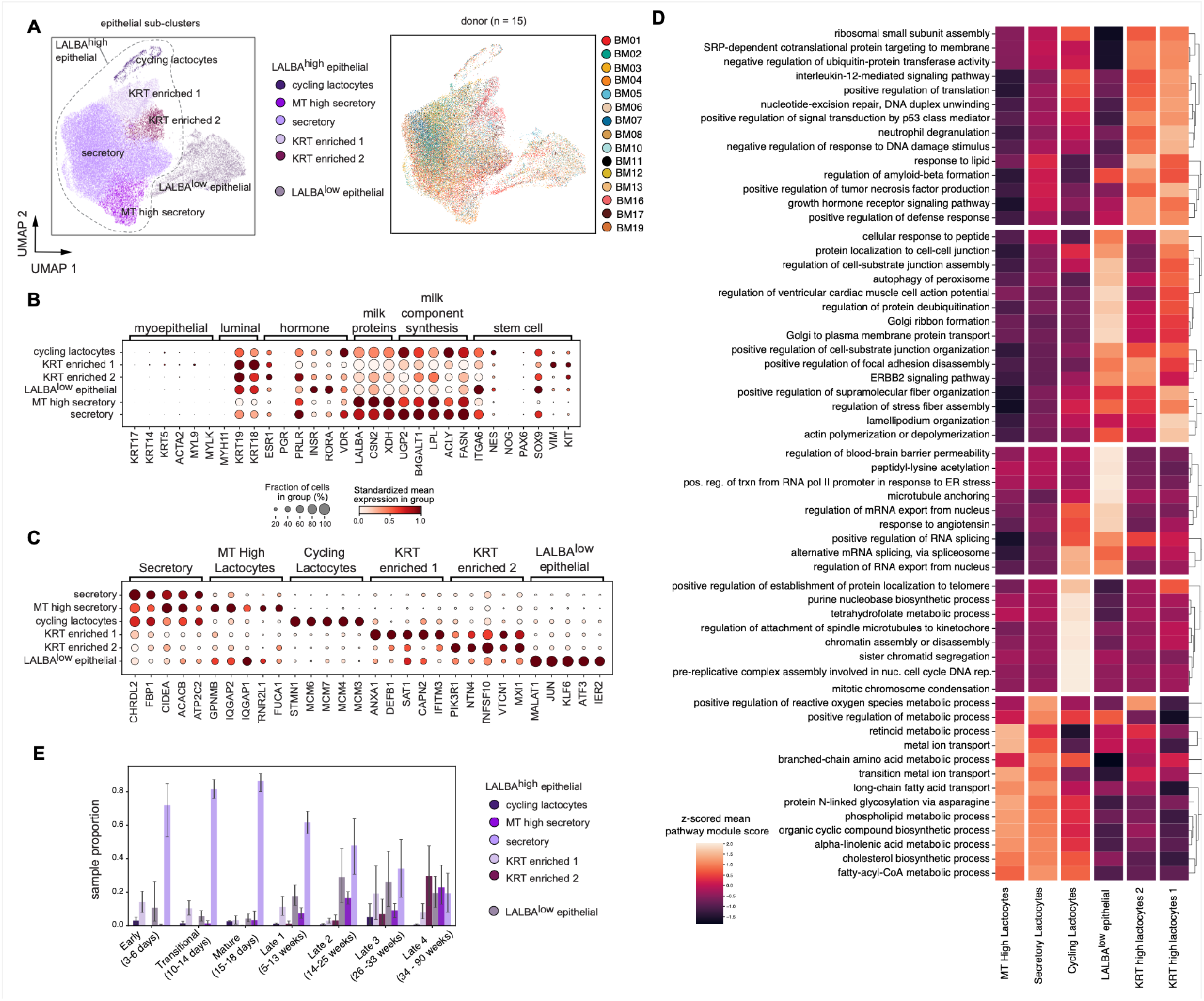
Sub-clustering analysis of epithelial cells reveals an increase in epithelial diversity over the course of lactation. **A**. UMAP visualization of epithelial cells colored by epithelial sub-cluster (left) or donor (right). **B**. Mean expression in cell subset standardized within genes (color) and percent of cells expression (dot size) of canonical mammary epithelial marker genes in each epithelial subgroup **C**. Mean expression in cell subset standardized within genes (color) and percent of cells expression (dot size) of marker genes for each epithelial subgroup identified by pseudobulk marker gene identification. **D**. Reduced top Enrichr results from the gene ontology biological processes 2021 database on the marker genes for each subgroup, colored by the mean gene set score for all genes in that pathway on cells in that subgroup, scaled by a z-score across subgroups. **E**. Proportions of each subgroup per sample, split by milk stage. Error bars show standard deviation.

In order to better understand the functions of each sub-cluster, we identified marker genes (Figure 4C) and performed enrichment analyses (Figure 4D). The largest sub-cluster of epithelial cells, secretory lactocytes, expressed the highest levels of secretory markers (*CHRDL2, CIDEA, ATP2C2*) and lipid and lactose synthesis genes (*FBP1, ACACB*). This cluster was also enriched for many pathways associated with metabolic processes, ion transport, and cholesterol biosynthesis. While there is significant heterogeneity within this large group of cells, this heterogeneity appeared as a continuum (see Methods, Supplemental Figure 4H and Supplemental Figure 5). The second largest sub-cluster, LALBA^low^ epithelial cells, was defined by expression of AP-1 transcription factor subunits (*JUN, ATF3, FOS*) as well as *MALAT1, KLF6* and *CLDN4*, genes involved in tight junction pathways.^67^ This sub-cluster was enriched for pathways related to microtubule and cellular organization (microtubule anchoring, actin polymerization or depolymerization), cell-cell junction assembly, protein transport via the golgi, and ERBB2 signaling pointing to an involvement in the establishment and maintenance of the cell-cell tight junctions which structurally support the alveolar structures in the lactating breast.

The cycling epithelial sub-cluster was defined by the expression of cell-cycle genes (*STMN1, TOP2A*) and was enriched for cell-cycle related processes as well as several metabolic processes, tetrahydrofolate metabolic process and purine nucleobase biosynthetic process. This sub-cluster is also composed entirely of cells whose cell-cycle score indicated they were in the G2M and S phases (Supplemental Figure 4C). The MT-high cluster was defined by similar gene expression to the secretory epithelial cells but with higher mitochondrial gene proportion (Supplemental Figure 4G). While mitochondrial RNA percentage is often used as a metric for dead or dying cells in scRNA-seq analysis, we maintained this cluster in the dataset because it met our very conservative threshold for mitochondrial RNA percentage, showed an interesting trend of increasing proportion over time, and may relate to altered metabolic activity in these cells.^68^

The KRT high lactocyte 1 cluster was defined by expression of cytoskeleton and structural genes (*S100A9, KRT15, KRT8, VIM*) as well as immune response genes (*ANXA1, DEB1, IFITM3, CD74, HLA-B*). This sub-cluster is enriched in genes in the actin polymerization or depolymerization pathway, positive regulation of defense response, positive regulation of translation pathways as well as several signaling pathways. The KRT high lactocyte 2 sub-cluster was enriched for similar pathways to the KRT high lactocyte 1 group, but this sub-cluster shares fewer high-scoring pathways with the LALBA^low^ lactocyte sub-cluster suggesting more of a supporting role in milk production.

Finally, we determined how these sub-clusters were changing in proportion as a function of lactation stage (Figure 4E). Globally, we found that the cellular composition of later lactational timepoints was more diverse as compared to earlier time points, where early time points are dominated by secretory epithelial cells. All sub-clusters, except the secretory and the cycling lactocytes, increase over the course of lactation. This may indicate that some degree of cellular specification is acquired over the course of lactation, potentially to meet changing demands on the maternal-infant dyad. For example, the increase in mitochondrial activity in the MT high sub-cluster, coupled with alterations in several metabolic pathways, may suggest that there are altered metabolic programs that support the high lactational demand and tissue turnover in later lactation.

### There were significant changes in gene expression over the course of lactation in the LALBA^low^ epithelial and secretory lactocyte sub-clusters

We found that both the fractional abundance and the overall epithelial diversity increased with time postpartum in hBM. So we next asked which genes and pathways also changed over the course of lactation in epithelial cells. To accomplish this, we performed differential expression with pseudo-bulk populations across time postpartum within each epithelial sub-cluster (see Methods). We found that there were many genes that were differentially expressed over time across all epithelial cells, including several that decrease over time such as *APP, KRT15*, and *FTH1*, and several that increase over lactational time, such as *LYZ* and *TCN1* (Figure 5A). Lysozyme, encoded by the transcript *LYZ*, one of the most abundant bioactive components of milk, has previously been shown to increase in later stages of lactation.^69^

**Figure 5:**
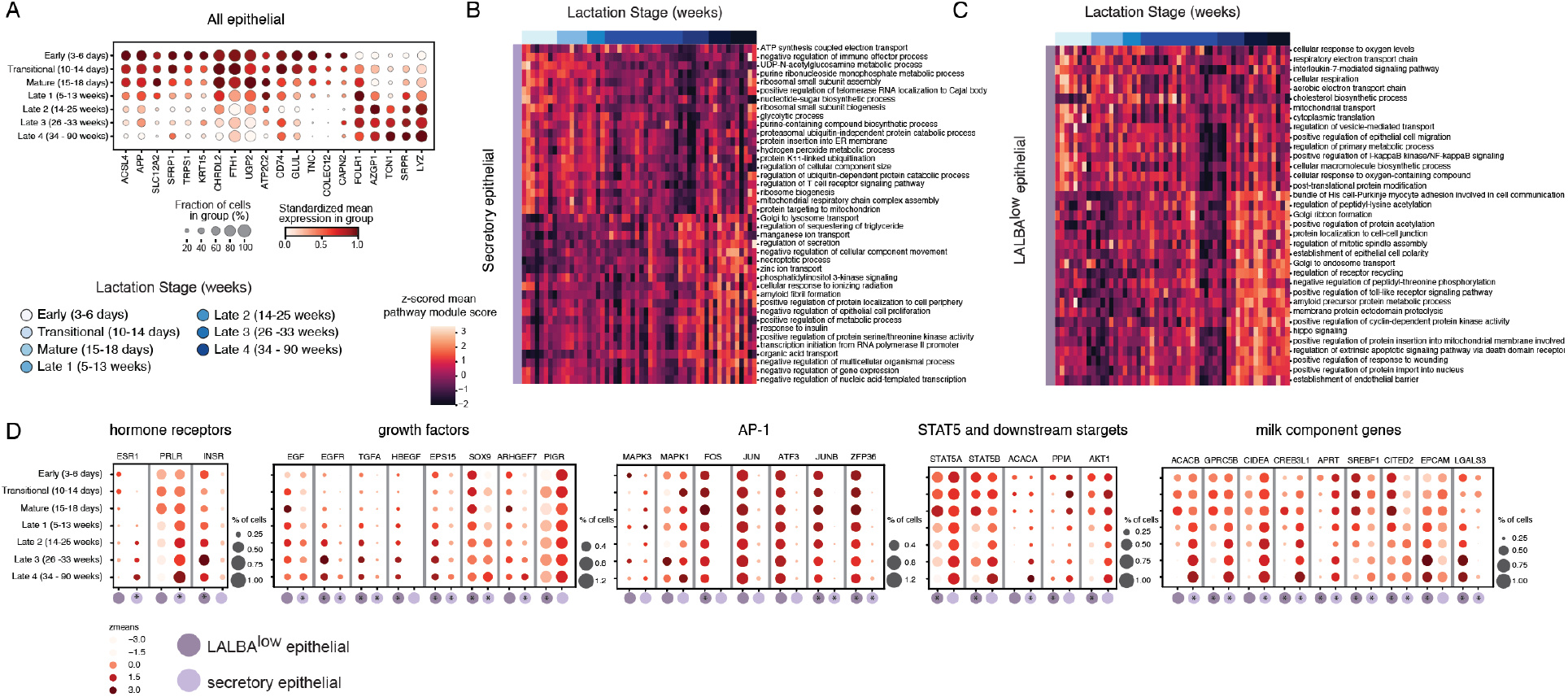
Transcriptional programs of luminal epithelial cells change over the course of lactation. A. Genes of interest changing over all epithelial clusters over the course of lactation, standardized expression over time. B. Reduced top enrichr GO biological process results on genes changing over time in secretory epithelial cluster and C. LALBA^low^ epithelial cluster heatmaps represent sample means of gene set scores of each pathway z-scored across samples, samples ordered by increasing time postpartum. D. hormone receptors, growth factor pathway components, AP-1 subunits, STAT5 and downstream targets, and milk component genes change with different dynamics in the LALBA^low^ epithelial and secretory lactocyte subclusters. Plots colored by mean expression of cells in each milk stage and time point z-scored across all time points and both subgroups.

Many genes were altered over the course of lactation that were unique to each identified epithelial sub-cluster, with the majority of differentially expressed genes identified in LALBA^low^ epithelial and secretory lactocyte sub-clusters (Supplemental Table 5A-B). Enrichment analyses of these differentially expressed genes (see Methods), identified both shared and distinct pathways that changed with time in both cell sub-clusters (Figure 5B, C and Supplemental Figure 6A). Several pathways change in expression over the course of lactation in secretory lactocytes, including a decrease in gluconeogenesis and oxidative phosphorylation over time, and an increase in the regulation of secretion and lipid metabolic processes (Figure 5B). The cholesterol biosynthesis pathway is enriched in both cell sub-clusters, but increases over time in secretory lactocytes and decreases over the course of lactation in the LALBA^low^ epithelial sub-cluster (Figure 5C). Additionally, over the course of lactation, pathway scores for TGF-beta signaling, chromatin remodeling factors, cytoskeletal transport, vesicle mediated transport, and apoptosis all increase in LALBA^low^ epithelial cells with time postpartum. Taken together, we identified many pathways that are differentially altered with lactation time in the major sub-clusters of epithelial cells.

In order to nominate key genes and factors that might be responsible for pathway-level changes in these two sub-clusters, we looked at the expression of key regulators that were differentially expressed with time postpartum, including those important for hormone signaling, growth factor signaling, AP-1 signaling, factors involved in STAT5 signaling, and several milk production component genes (Figure 5D)^70–72^. We found that the expression of several hormone receptor genes changed in opposite directions with time in the LALBA^low^ epithelial and secretory lactocyte sub-clusters, where estrogen receptor (*ESR1*) and prolactin receptor (*PRLR*) increased in secretory lactocytes and decreased in LALBA^low^ epithelial cells.^13,14^ Insulin receptor (*INSR*) increased in just LALBA^low^ epithelial cells. Given that these receptors are crucial to orchestrating the functions and tissue structure of the lactating mammary gland, our data suggests that these two sub-clusters may differentially contribute to these functions over time in a hormonally-regulated manner.

## DISCUSSION

In this study, we used scRNA-seq to provide the first in-depth characterization of the transcriptional changes over the course of lactation in hBM in a single cell manner. Our cohort represented a wide range of experiences of maternal-infant dyads that allowed us to determine how cellular content varied over the course of lactation, and which maternal and infant factors (metadata features) were correlated with hBM cellular content, how cells changed their transcriptomes longitudinally, and what the full depth of cellular diversity was over each lactation stage.

We found that the majority of immune cells in our data were macrophages and that adaptive immune cells, including T cells and B cells, were only a small fraction of the total recovered cells from hBM. Our top-level clustering revealed two major populations of macropahges, both enriched for canonical macrophage markers like CD68. We found that our CSN1S1^+^ macrophage cluster was enriched for several milk production transcripts, like *CSN*. These could be present in this population as “passenger” transcripts that originate from engulfed apoptotic bodies or these may be functionally important given previously defined ductal associated macrophages express similar milk-related transcripts.^19,73^ We also identified several sub-clusters of macrophages, and our GO enrichment and module scoring analyses suggests that these may be more tolerogenic in nature. Previous reports in mice have found extensive diversity in mammary duct macrophages, and have found that these cells alter their transcriptomes significantly over reproductive cycles.^19^ This, coupled with work in the context of breast cancer and pan tissue analyses, suggests that the full functional diversity of macrophages in the human breast has yet to be fully characterized.^74^ Future work should seek to better understand the factors that promote tolerogenic functions of macrophages during lactation, whether its tissue specific or milk specific factors, and what secreted factors from macrophages might support healthy mammary gland functions. The association of our macrophage GPNMB^+^ cluster with formula supplementation was interesting, but our cohort was not powered to investigate potential mechanisms. Future work should seek to understand how formula might alter cellular composition in hBM, and whether this could impact the functions of hBM-derived macrophages.

Through sub-clustering analyses on epithelial cells, we identified two major populations of epithelial cells (LALBA^high^ and LALBA^low^) as well as several sub-clusters of LALBA^high^ epithelial cells (cycling lactocytes, KRT enriched 1, KRT enriched 2, secretory, MT high secretory). These agree with previous reports, underscoring the functional diversity of these cells and their difference as compared to breast tissue.^16^ Our data suggests that LALBA^low^ epithelial cells may provide more structural support during later lactation stages, while LALBA^high^ epithelial cells and its associated sub-clusters may produce more milk components. Consistent with previous work, we also did not see cells expressing genes expected from myoepithelial, basal or stem cells.^16,29^

Unlike previous reports, our data provided a unique opportunity to determine how cell types change in both composition and function over the full course of lactation, and if these changes are associated with maternal-infant metadata. Our data suggests that milk is dynamic over the full course of lactation, with immune cells expanding and contracting within each sample over time. Previous reports have well-defined infiltration of CD45^+^ cells in response to mastitis and other infections, and have characterized extensively the features of immune cells by canonical makers in the context of pre-term birth or infection^3,18,25,60,75^. These studies predominantly relied on flow cytometry, and here, we were able to use scRNA-seq to in depth characterize alterations in cellular composition with less potential bias. Given our limited sample processing (e.g. no staining or sorting), we may have also recovered more macrophages than previous studies. To our knowledge, our study is also the first to correlate maternal-infant dyad metadata with cell proportions over the full course of lactation. We found that the proportion of LALBA^low^ epithelial cells and B cells were associated with time postpartum using generalized additive models; however, we acknowledge that the overall frequency of B cells in our final dataset was low and precluded more in-depth analyses. Given that B cells are critical to the production of antibodies and these in turn shape early immune system development, future studies should seek to compare B cell repertoires from hBM and in circulation to better delineate how antibodies are transferred to hBM and the importance of these cells in the lactating mammary gland.

In addition to being correlated with time postpartum, the proportional abundance of LALBA^low^ epithelial cells were positively associated with two external factors: daycare attendance and hormonal birth control usage. The effect of these variables is challenging to disentangle in our dataset, but our results suggest that future work should specifically seek to understand how external perturbations and behaviors, potentially including increased pumping frequency and circulating hormone levels, impact the mammary gland specifically during later stages of lactation. Our differential expression results identifying key growth factors and hormone receptors, like ESR1 and INSR, that changed in expression over time in these cells suggests that these may be hormonally regulated and emerge as important structural cells in later stages of lactation.

At the gene level, bulk transcriptomic studies have shown transcriptional changes between colostrum, transitional and mature milk in pathways presumed to originate from epithelial cells, indicating that insulin signaling, lactose synthesis, and fatty acid synthesis pathways increase during these early stages of lactation.^26^ Only a few transcriptional studies have characterized the gene expression changes during later stages of lactation before involution. While previous studies show higher expression levels of *PRLR, STAT5A*, and milk protein and lipid synthesis genes during lactation when compared to colostrum or involution, bulk longitudinal studies have not had the resolution to describe the changes in cells co-expressing these genes.^12,27^ Additionally, more milk components are transferred from the blood to the milk via tight junctions at later time points in lactation and fewer components are synthesized in the lactocytes themselves.^76^

We provide, in great detail, the epithelial cell sub-clusters in which key genes are changing across both time and many donors, allowing us to gain insights into potential alterations in milk transport, synthesis, and production. The LALBA^low^ epithelial cell cluster, whose marker genes are enriched for genes involved in tight junctions, increases in abundance over the course of lactation while we see a decrease in the proportional abundance of the secretory lactocyte sub-cluster whose core enriched functions involve milk component synthesis and secretion. We also see a decrease in milk component synthesis related genes (UGP2, CHRDL2) (Figure 5A) and a decrease in the GO terms gluconeogenesis, hexose biosynthetic process, glucose metabolic process over time in both clusters (Figure S6). This might suggest a decrease in transcription of milk component related genes over the course of lactation. Previous studies have shown a linear decrease in overall protein concentration in milk over the course of lactation as well as decrease in concentrations of proteins involved in lactose and HMO synthesis.^76–78^ In addition, due to our long follow up study, we were able to capture late stages of mature milk (late 2-4), when usually complimentary food are presented to babies, and milk demand and production decreased over time. Increased cellular specialization and altered abundance of epithelial sub-clusters that we describe may provide mechanistic insights into changes in the maintenance of milk secretion over the course of lactation. Future work should specifically seek to understand how this relates to milk component production as synthesis in the mammary gland, transport from maternal serum, or milk volume production.

Hormones in hBM serve both as regulators of the mammary gland itself as well as bioactive components passed to the infant. Lactogenesis and the initiation of lactation at the end of pregnancy are tightly hormonally regulated by a drop in serum progesterone allowing prolactin signaling to initiate lactation.^13^ Milk component synthesis and secretion during peak lactation have also been shown to be regulated more locally in the mammary gland by milk removal, autocrine hormone signaling, and in the lactocytes themselves.^79^ Prolactin receptor (*PRLR*) is known to be involved in many aspects of the continuation of lactation,^14,80^ and prolactin concentration in breast milk decreases over the course of lactation.^81^ We found that pathways downstream of several hormone receptors, including prolactin signaling, estrogen signaling, and human growth factor signaling, were enriched in the marker genes of the LALBA^high^ epithelial cells, indicating that these cells are likely directly hormonally regulated.

Interestingly, the LALBA^low^ epithelial and secretory epithelial cell sub-clusters showed opposite changes in hormone receptor expression over the course of lactation (Figure 5D), pointing to a possible regulatory mechanism of these synthesis and transport changes vis a vis a division of labor between cell types potentially over the course of lactation. *STAT5A* is a core lactational gene that is involved in proliferation, cell survival, and milk component synthesis.^70,80,82–86^ Interestingly, we observe decreases in *STAT5a/b* expression and downstream targets such as *AKT1*^72^, *ACACA* (a gene involved in fatty acid synthesis), and *CSN2* (the gene encoding beta-casein) over the course of lactation. We also found a decrease in the GO terms cellular macromolecule biosynthetic process and cholesterol biosynthetic process in LALBA^low^ epithelial cells over the course of lactation, all of which are related to milk component synthesis^82,85–87^. In secretory epithelial cells, expression of *PRLR* increases with time postpartum and some increase in *JAK2* expression and *STAT5A* expression are also observed as well as target *ACACA* in this cell subset. Taken together, our data suggests that these two groups of epithelial cells may shift in their responsiveness to prolactin and prolactin-regulated *STAT5* pathways over the course of lactation. This shift could explain other differential functions of these cell sub-clusters over the course of lactation if, for example, the LALBA^low^ epithelial cells become more responsible for milk component transport over the course of lactation and increase their prolactin and JAK2/STAT5 regulated milk component synthesis. We see similar alterations in the dynamic expression of several growth factors that regulate milk production and secretion, ^88^ like *EGF*. Further studies should investigate this division of cellular labor and consider the direction of this regulation and how it might be leveraged therapeutically to potentially aid in milk production.

## CONCLUSION

Human breast milk is a dynamic living fluid that contains millions of cells. Here, we used scRNA-seq to characterize the transcriptomes of single cells from hBM across lactational time. We confirm that the majority of cells in human breast milk are epithelial cells, and specifically lactocytes, and that cell type frequencies are dynamic over the course of lactation. Analysis of lactocytes reveals a continuum of cell states characterized by subtle transcriptional changes in hormone, growth factor, and milk production related pathways, that occurs over the course of lactation. These results point to changing populations of milk component-producing epithelial cells whose activities over the course of lactation may be hormonally regulated. We also identify several sub-clusters of macrophages in hBM that are enriched for tolerogenic functions. Taken together, our data provide the first detailed longitudinal study of breast milk cells with single-cell resolution. Further understanding of cells over the course of lactation, including B cells, macrophages, and LALBA^low^ epithelial cells, will build knowledge of the role of breast milk in infant development by identifying: (i) cells that are transferred to infant gut, (ii) the molecules they produce that are important for gut ^6,89^ and immune system development, and (iii) the nutrients supplied in hBM.

Our description of the cellular components of breast milk over the course of lactation, and their association with maternal-infant dyad metadata, has the potential to provide insights into mechanisms of milk-component production and regulation, as well as variability between individuals^1^. Improved understanding of pathways and activities of breast milk producing cells will add to the understanding of lactation health and could provide baseline information for studies of adverse lactation outcomes. Lastly, studies of long term lactation, such as ours, will aid in establishing eligibility criteria for milk bank donation potentially allowing donors to contribute milk after the typical one year postpartum limit.^90^

## ACKNOWLEDGEMENTS

We thank the study participants and their families for enabling this research, Nancy Tran and other members of the Shalek and Berger labs for thoughtful discussions and feedback. We thank the Single Cell Portal, Terra, and Cumulus teams at the Broad Institute for support on data processing pipelines and data sharing. This work was supported in part by the Koch Institute Support (core) NIH Grant P30-CA14051. BAG was supported by NRSA postdoctoral fellowship (F32-AI136459). AKS was supported, in part, by the Beckman Young Investigator Program, a Sloan Fellowship in Chemistry, and MIT (Charles E. Reed Faculty Initiative). SKN was supported by National Science Foundation Graduate Research Fellowship (1122374). BEM by MIT-GSK Gertrude B. Elion Postdoctoral Fellowship. MEM by Columbia University Office of the Provost grants for junior faculty who contribute to the diversity goals of the University. YGM was supported by Weizmann Institute of Science -National Postdoctoral Award Program for Advancing Women in Science, the International Society for Research In Human Milk and Lactation (ISRHML) Trainee Bridge Fund, and of the Human Frontier Science Program (HFSP).

## AUTHOR CONTRIBUTIONS

BAG, SKN, and AKS Conceived of the stud y. BAG designed the study. BAG, SKN, KK, RSD, and BEM optimized study protocol. BAG, SKN, KK, collected and processed samples. BAG, SKN, PG, TJKH, MRR, KK performed single-cell sequencing experiments. BAG, SKN, PG, MRR analyzed data under supervision of MEM, AKS, and BB. YGM and NA assisted in interpretation of data. BAG and SKN wrote the original draft. AKS, YGM, MEM, BEM, NA contributed to the manuscript and all authors provided comments. AKS and BB acquired funding and provided resources.

